# Neuromodulatory selection of motor neuron recruitment patterns in a visuomotor behavior increases speed

**DOI:** 10.1101/683649

**Authors:** Urvashi Jha, Vatsala Thirumalai

**Affiliations:** National Centre for Biological Sciences, Tata Institute of Fundamental Research; Bangalore; Karnataka-560065; India; SASTRA Deemed University; School of Chemical and Biotechnology; Thanjavur; Tamil Nadu-613401; India

**Keywords:** Dopamine, optomotor response, zebrafish, spinal cord, excitability, D1-like receptor

## Abstract

Animals generate locomotion at different speeds to suit their behavioral needs. Spinal circuits generate locomotion at these varying speeds by sequential activation of different spinal interneurons and motor neurons. Larval zebrafish can generate slow swims for prey capture and exploration by activation of secondary motor neurons and much faster and vigorous swims during escapes and struggles via the additional activation of primary motor neurons. Neuromodulators are known to alter motor output of spinal circuits yet their precise role in speed regulation is not understood well. Here, in the context of optomotor response (OMR), an innate, evoked locomotor behavior, we show that dopamine (DA) provides an additional layer to regulation of swim speed in larval zebrafish. Activation of D1-like receptors increases swim speed during OMR in free-swimming larvae. By analysing tail bend kinematics in head-restrained larvae, we show that the increase in speed is actuated by larger tail bends. Whole cell patch clamp recordings from motor neurons reveal that during OMR, typically only secondary motor neurons are active while primary motor neurons are quiescent. Activation of D1-like receptors increases motor drive from secondary motor neurons by decreasing spike threshold and latency. In addition, D1-like receptor activation enhances excitability and recruits quiescent primary motor neurons. Our findings provide an example of neuromodulatory reconfiguration of spinal motor neuron speed modules such that members are selectively recruited and motor drive is increased to effect changes in locomotor speed.

**Highlights:**

- Zebrafish larvae generate swims of increased speed during optomotor response when D1-like receptors are activated.
- D1-like receptor activation increases the extent of tail bending during forward swims and turns resulting in increased swim speed.
- Neuromodulation via D1-like receptors increases motor drive by enhancing excitability of ‘slow’ motor neurons. In addition, D1-like receptor activation recruits quiescent ‘fast’ motor neurons to increase swim speed.
- This demonstrates neuromodulatory selection of motor neurons belonging to different ‘speed’ modules to alter swimming behavior.

**Graphical abstract:** 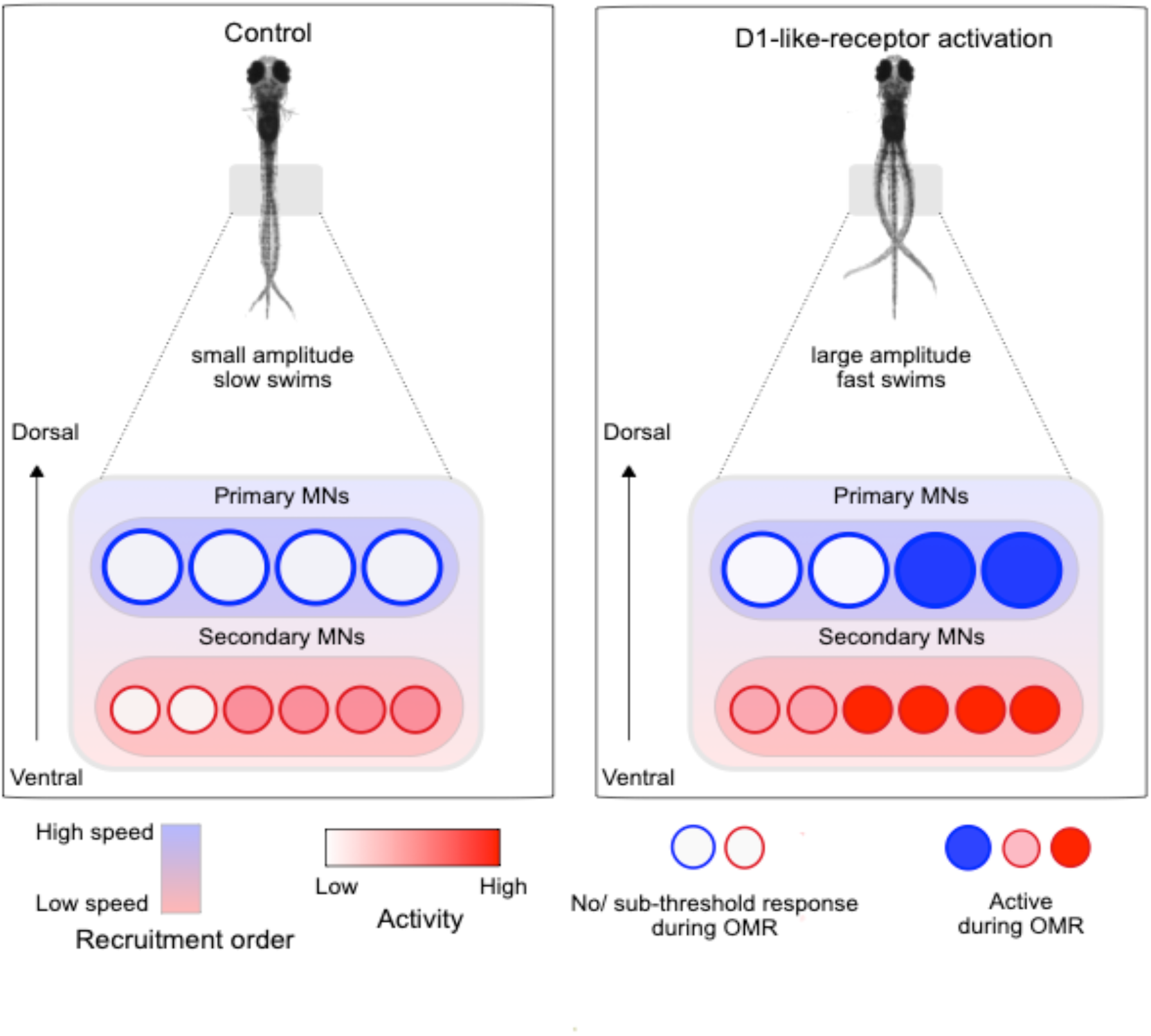

## Introduction

Survival for most animals depends on their ability to adapt speed, gait and direction of movement quickly in response to sensory stimuli. In vertebrates, these processes are controlled by the concerted activity of distributed circuits in the brain and the spinal cord. In particular, the regulation of speed by spinal locomotor circuits has received much attention recently. The ability to genetically label specific neuronal populations and to target them for imaging or electrophysiology has allowed us to identify recruitment patterns of interneurons and motor neurons at different speeds. Studies in mice and zebrafish have identified subtypes of spinal interneurons and motor neurons that are selectively recruited at different speeds of locomotion [1–8]. These interneurons and motor neurons make selective synaptic connections among them and are thought to constitute ‘slow’ or ‘fast’ network modules, that are sequentially recruited as the locomotor speed increases [9, 10]. Thus, behaviors such as escapes, which involve fast swimming, are generated by the fast module, and spontaneous swimming behaviors, occurring at much lower swim frequencies are controlled by the slow module [11].

Although this modular view of spinal network operation is widely acknowledged [12, 13], we also know that the spinal network is the target of a variety of neuromodulators [14–17]. Neuromodulators have been shown to fuse independent circuits and to construct ‘de novo’ circuits from cells belonging to disparate circuits [18, 19]. We wished to understand if swim speed regulation in zebrafish larvae is subject to neuromodulation and if yes, whether the rules for modular recruitment are altered by neuromodulators.

Dopamine has long been known to be a key neuromodulator of motor circuits [17, 20] and is critical for activation, maturation and modulation of locomotor patterns across invertebrates and vertebrates [21–26]. In zebrafish, dopaminergic input from the diencephalon to the spinal cord develops early [27] and is known to regulate developmental maturation of swim motor patterns [24]. Nevertheless, the role of spinal dopamine in regulating speed of locomotion has thus far not been investigated in any vertebrate.

We chose to study speed regulation and its dopaminergic modulation, during the optomotor response (OMR), an innate behaviour evoked by whole-field visual motion, to which zebrafish larvae respond by turning towards and moving in the direction of perceived motion. By adjusting their locomotor speed to match the speed of the perceived motion, larvae try to maintain a stable position with respect to their surroundings [28, 29]. However, the spinal mechanisms by which speed is controlled during OMR have thus far not been investigated.

We used a combination of behavioral monitoring in free-swimming larvae, kinematic characterization in head-restrained larvae and whole-cell patch clamping in immobilized larvae to ask how speed is controlled during OMR and how it is modulated at the neuronal level. Here, we show that dopamine acting via D1-like receptors (D1-like-R) increases the speed of swimming during OMR by increasing the extent of tail bending. This in turn is caused by two factors: (1) increased firing of action potentials in motor neurons of the slow module, typically recruited during OMR and (2) novel recruitment of motor neurons belonging to the fast module, not typically active during OMR. Thus, dopamine selectively activates and recruits neurons from disparate spinal modules to increase speed. Our results suggest that activity and recruitment pattern of neurons in these circuit modules are not only defined by their cellular properties and synaptic connectivity. Neuromodulatory inputs can dramatically alter the dynamics of these modules by recruiting neurons that would have otherwise not participated under a given context.

## Results

### D1-like-R activation increases swim speed during OMR

OMR was evoked in freely swimming zebrafish larvae between 6-7 days post fertilization (dpf) by presenting square-wave gratings moving in a clockwise direction from below (Figure 1A, Video S1). Zebrafish larvae respond by following the direction of the stimulus and matching their swim speed with that of the moving grating [29]. Larval centroid was tracked to extract kinematic parameters. Zebrafish larvae swim intermittently in a beat-and-glide pattern where brief periods of discrete tail oscillations (bout) are followed by a period of no tail movement [30]. We further segmented tracked data to identify individual swim bouts (Figure 1B).

**Figure 1.**
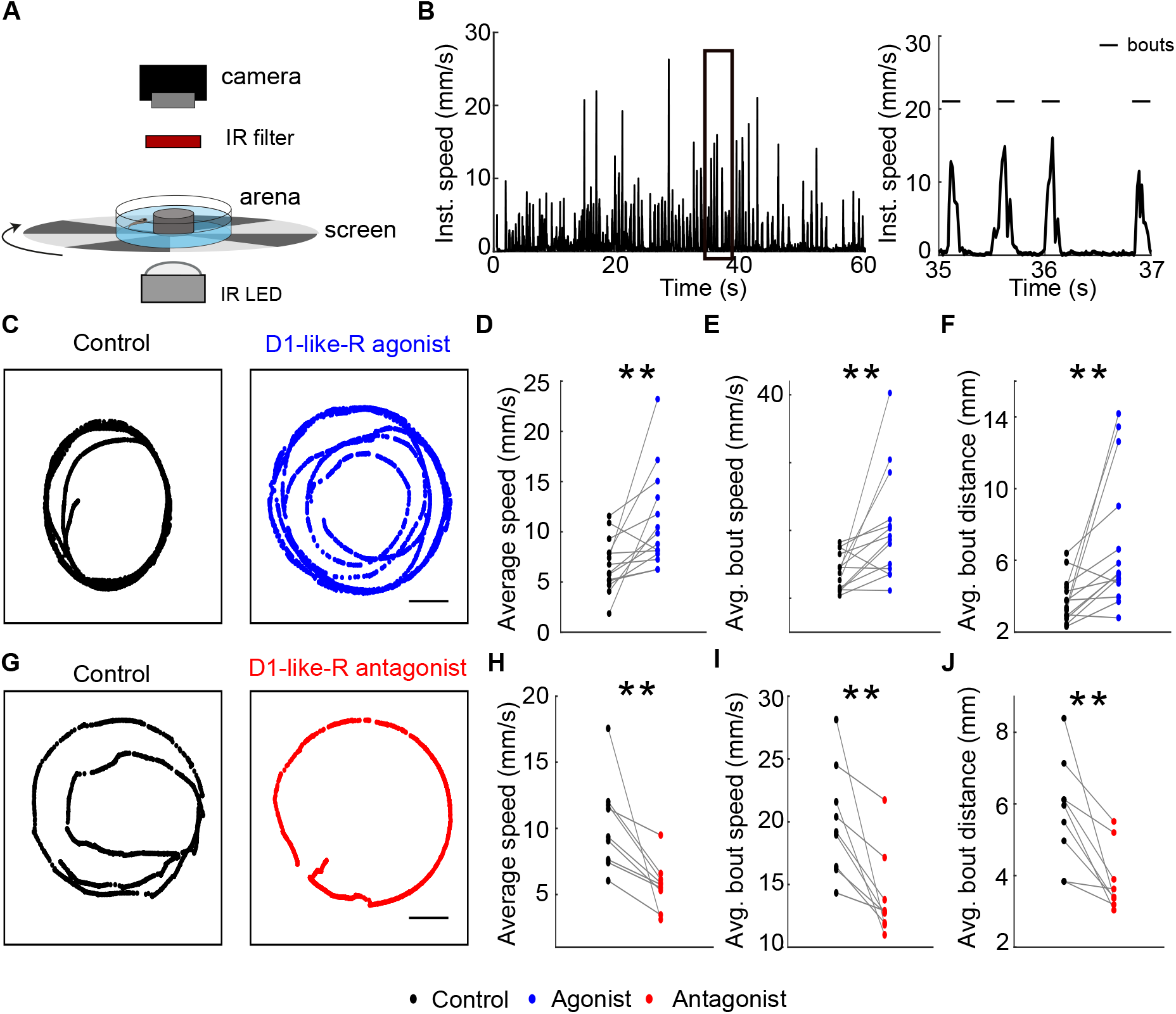
D1-like-R activation increases swim speed during OMR in free-swimming zebrafish larvae. **(A)** Schematic of experimental set up. OMR was evoked in freely swimming zebrafish larvae (6-7 dpf) by presenting radial gratings moving in a clockwise direction on a screen below the arena. Zebrafish larvae respond to the stimulus by swimming in the direction of moving grating. Videos were acquired from above and larval centroid was tracked. **(B)** *Left*, instantaneous speed of a representative larva during a trial. *Right*, zoomed view of region marked with the black rectangle showing characteristic intermittent swimming pattern. Brief period of activity represents individual swim bouts (highlighted by line on top) and is followed by a period of no tail movement. **(C)** Overlaid tracked centroid for a trial duration of 60 seconds showing trajectory of a representative larva before (Control) and after bath application of 20 μM D1-like-R agonist, SKF-38393. Scale bar represents 10 mm. **(D-F)** Paired plots for average speed, average bout speed and average bout distance respectively, before (black) and after (blue) application of D1-like-R agonist. **(G)** Overlaid tracked centroid of a representative larva for a trial duration of 60 seconds before (Control) and after bath application of 20 μM D1-like-R antagonist, SCH-23390. Scale bar represents 10 mm. **(H-J)** Paired plots for average speed, average bout speed and average bout distance respectively, before (black) and after (red) application of D1-like-R antagonist. N_SKF-38393_=14 larvae, N_SCH-23390_=9 larvae, Wilcoxon signed rank test; ** p<0.001. See also Video S1.

We compared average swim speed (including both bouts and inter-bout quiescent periods) during OMR before and after bath application of 20 µM D1-like-R agonist, SKF-38393. Larvae swam with higher average swim speed post D1-like-R activation (Figure 1C, D, Video S1). We then asked if D1-like-R activation increases speeds achieved during bouts and compared average bout speed before and after bath application of D1-like-R agonist. Average bout speed increased (Figure 1E) resulting in greater distance travelled per bout (Figure 1F) post application of D1-like-R agonist.

To further confirm that the increase in swim speed is a result of D1-like-R activation and not an artefact of repeated stimulus presentation, we also compared these parameters before and after bath application of 20 µM D1-like-R antagonist, SCH-23390. Average swim speed decreased post application of D1-like-R antagonist (Figure 1G, H). Consistently, a decrease in average bout speed (Figure 1I) and average distance travelled per bout (Figure 1J) were observed post application of D1-like-R antagonist.

D1-like-R signaling is associated with increased spontaneous locomotor activity in larval zebrafish [24, 31, 32]. Similar to spontaneous swimming, we find D1-like-R signaling to have a stimulatory effect on the motor output during OMR. Importantly, these results indicate that locomotory speed during an innate and reflexive behavior is subject to neuromodulation via D1-like-R activation.

### D1-like-R signaling controls swim speed by modulating tail beat amplitude

We sought to further understand which kinematic variables are altered by D1-like-R signaling to bring about the increase in bout speed we observe. For characterization of kinematic variables, we evoked OMR in a head-restrained preparation which allows for a more accurate and detailed tracking of tail kinematics.

In response to caudal-to-rostral moving gratings, partially restrained larvae reliably performed forward OMR characterized by mostly symmetrical, alternating left-right tail beats (Figure 2A, Video S2). Tail-tip position was tracked to extract tail beat amplitudes, tail beat frequency and duration of bouts (Figure 2B). As DA has been shown to modulate swim initiation in larval zebrafish [33], we also quantified the number of swim bouts initiated.

**Figure 2.**
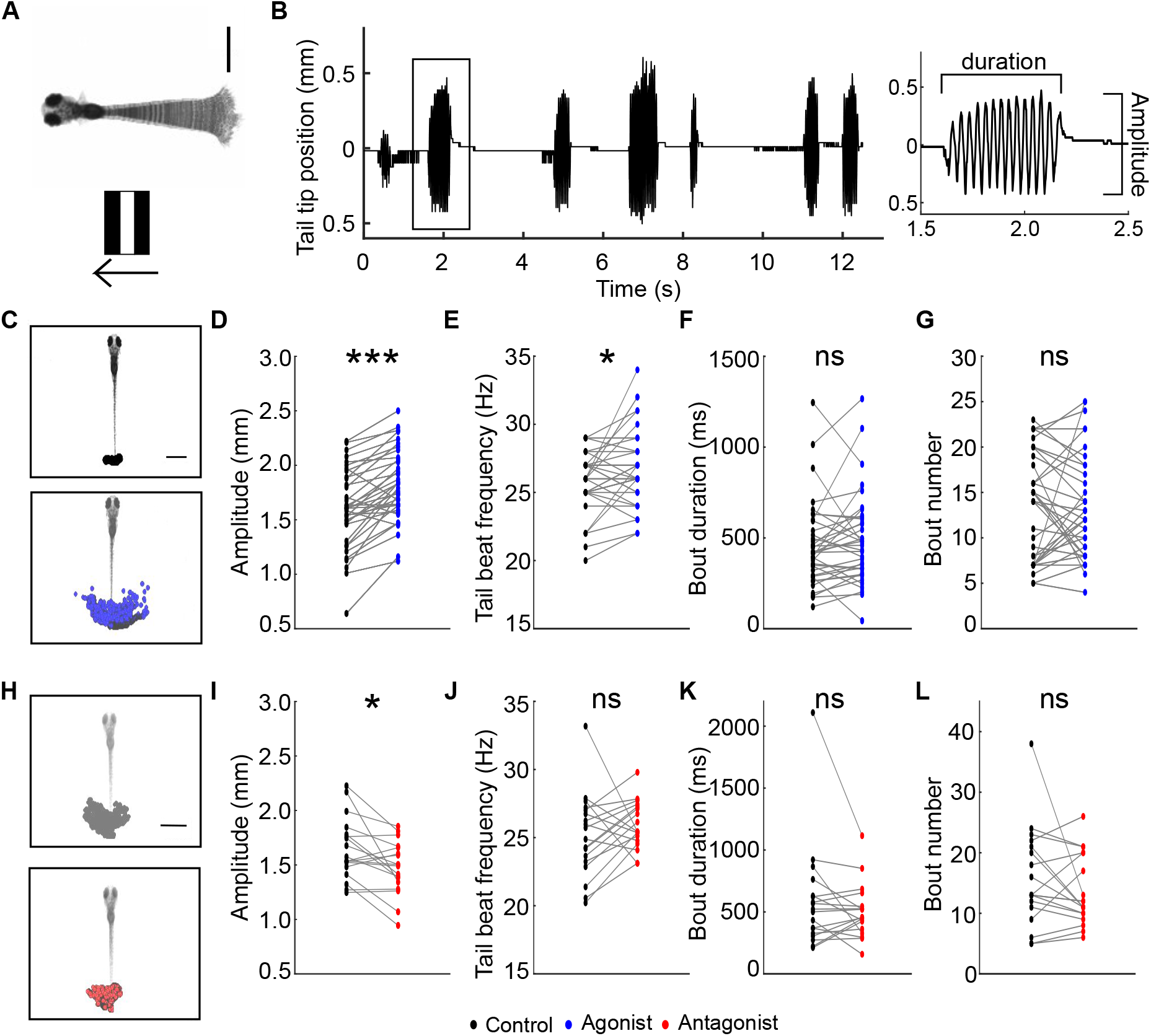
D1-like-R signaling modulates tail beat amplitude during forward OMR in head restrained larvae. **(A)** Schematic representation of experimental set-up. Z-projection of a swim bout showing characteristic alternating left-right tail beats evoked in a head-restrained larval zebrafish (top) in response to caudal-to-rostral moving gratings (bottom). Tail-tip position was tracked for measurement of kinematic variables. (**B)** *Left*, representative tracked tail-position for a trial. *Right*, zoomed view of highlighted bout showing kinematic variables used for behavioral quantification. **(C)** Overlaid tracked tail-tip position for a representative trial before (top) and after (bottom) application of 100 μM D1-like-R agonist, SKF-38393. Summary data of **(D)** average tail beat amplitude, **(E)** average tail beat frequency, **(F)** average bout duration and **(G)** number of bouts initiated before (black) and after (blue) bath application of D1-like-R agonist. **(H)** Overlaid tracked tail-tip position for a representative trial before (top) and after (bottom) application of 100 μM D1-like-R antagonist, SCH-23390. Summary data of **(I)** average tail beat amplitude, **(J)** average tail beat frequency, **(K)** average bout duration and **(L)** number of bouts initiated before (black) and after (red) bath application of D1-like-R antagonist. Scale bar represents 1mm. N_SKF-38393_=37 larvae, N_SCH-23390_=18 larvae, Wilcoxon signed rank test; *** p<0.00001, * p<0.05, ns: not significant. See also video S2.

Activation of D1-like-R increased mean tail beat amplitude and tail beat frequency of forward swims (Figure 2C-E, Video S2). However, no difference in duration of bouts or number of bouts initiated was observed post application of 100 µM D1-like-R agonist, SKF-38393 (Figure 2F, G). In contrast, application of 100 µM D1-like-R antagonist, SCH-23390, decreased mean tail beat amplitude of forward locomotion (Figure 2H, I). Tail beat frequency, bout duration and number of swim bouts initiated did not change significantly (Figure 2J-L).

Modulation of tail beat amplitude is a key factor contributing to control of swim speed [34, 35]. Thus, by modulating tail beat amplitude D1-like-R activation results in faster swim bouts during forward OMR.

### Activation of D1-like-R results in larger tail bend angle during turns

Free swimming OMR behavioural assay required larvae to swim both in a straight trajectory as well as make routine turns (Video S1). Since we observed an increase in both average swim speed and average bout speed after D1-like-R activation during free-swimming OMR (Figure 1), we surmised that D1-like-R activation will likely affect tail bend kinematics during routine turns also. To investigate the role of D1-like-R signaling in modulation of turning behavior, we presented leftward and rightward moving gratings to head-restrained larvae (Figure 3A). Larvae responded to the stimuli with asymmetric tail deflections in the direction of grating movement (Figure 3A, Video S3). Tail-tip position was tracked and maximum tail bend angle was measured before and after application of 100 µM D1-like-R agonist and antagonist (Figure 3B).

**Figure 3.**
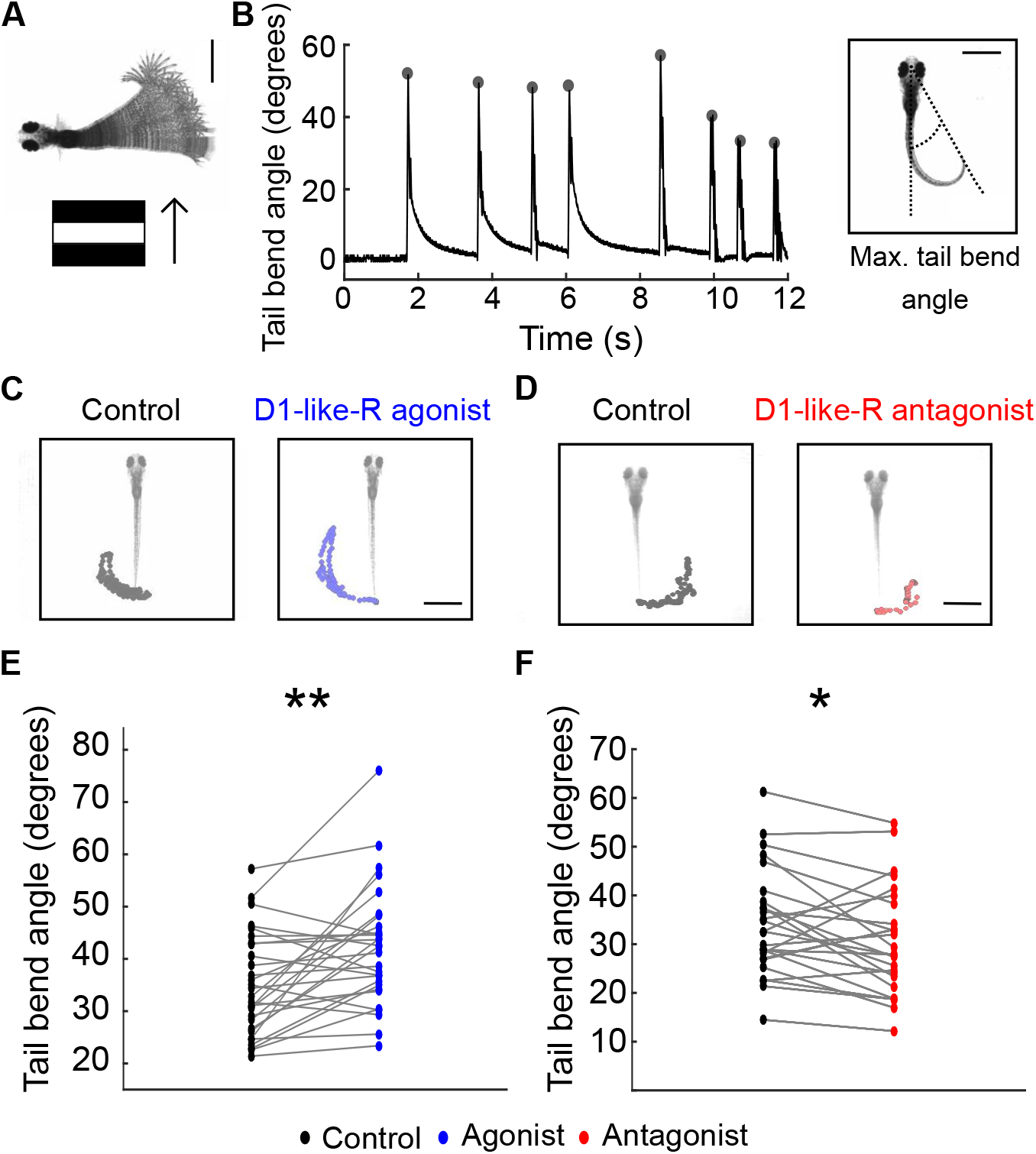
D1-like-R activation increases tail bend angle during turns. **(A)** Schematic representation of experimental set-up showing a turn characterized by asymmetric tail bend about the midline (top) evoked using sideward moving vertical bars (bottom) in a head-restrained zebrafish larva. Tail-tip position was tracked for measurement of tail bend angle. **(B)** *Left*, tail bend angle for a representative trial. Maximum tail bend angle for each turn is highlighted with a solid circle. *Right*, snapshot of a turn showing references used for calculation of tail bend angle. **(C)** Overlaid tracked tail-tip position during a trial before (left) and after (right) bath application of 100 μM D1-like-R agonist, SKF-38393. **(D)** Overlaid tracked tail-tip position during a trial before (left) and after (right) bath application of 100 μM D1-like-R antagonist, SCH-23390. **(E-F)** Summary data of average tail bend angle before and after application of D1-like-R agonist and antagonist. Scale bar represents 1mm. N_SKF-38393_=29 larvae, N_SCH-23390_=26 larvae, Wilcoxon signed-rank test; **p<0.001, *p<0.05. See also video S3.

Larvae performed turns with larger tail bend angles post application of D1-like-R agonist (Figure 3C, E, Video S3). In contrast, application of the D1-like-R antagonist showed a decrease in the magnitude of the tail bend angle (Figure 3D, F).

Taken together, these results indicate that D1-like-R activation modulates the extent of tail bends during both forward locomotion and turns resulting in greater displacement.

### Activation of D1-like-R intensifies drive from ‘slow’ motor neurons

D1-like-R mediated increase in swim speed could result from an increase in the firing of active motor neurons and/or by the recruitment of new cells to the active pool [1]. Zebrafish axial motor neurons can be broadly divided into two classes: primary and secondary [36]. Early born, dorsally located primary motor neurons innervate fast muscle fibres and are active during fast swimming, struggle and escape responses swimming whereas later born, ventral secondary motor neurons are active during slower swimming [1, 37]. Besides anatomical position and axonal innervation pattern, the two classes of motor neurons can also be clearly distinguished based on their cellular properties and response to injected current [38] (Table S1, Figure S1). As secondary motor neurons are active during low-frequency swimming such as those observed in OMR, we first asked if they are modulated by D1-like-R. We obtained whole-cell patch clamp recordings from dorsal secondary motor neurons while presenting moving gratings from below to evoke forward OMR (Figure 4A). 7/9 recorded dorsal secondary motor neurons fired bursts of action potentials in response to moving grating presentation (Figure 4B). An increase in the number of action potentials generated during moving gratings presentation was observed post application of 20 µM D1-like-R agonist (Figure 4C, G). Interestingly, we also observed motor neurons otherwise quiescent during visual stimulus presentation (2/9), fire action potentials post application of D1-like-R agonist (Figure 4D, E). D1-like-R agonist mediated effects were antagonized in 4/4 secondary motor neurons post subsequent application of D1-like-R antagonist (Figure 4F, G). Number of action potentials spontaneously generated during stationary grating presentation did not differ significantly before and after application of D1-like-R agonist (Figure S2A).

**Figure 4.**
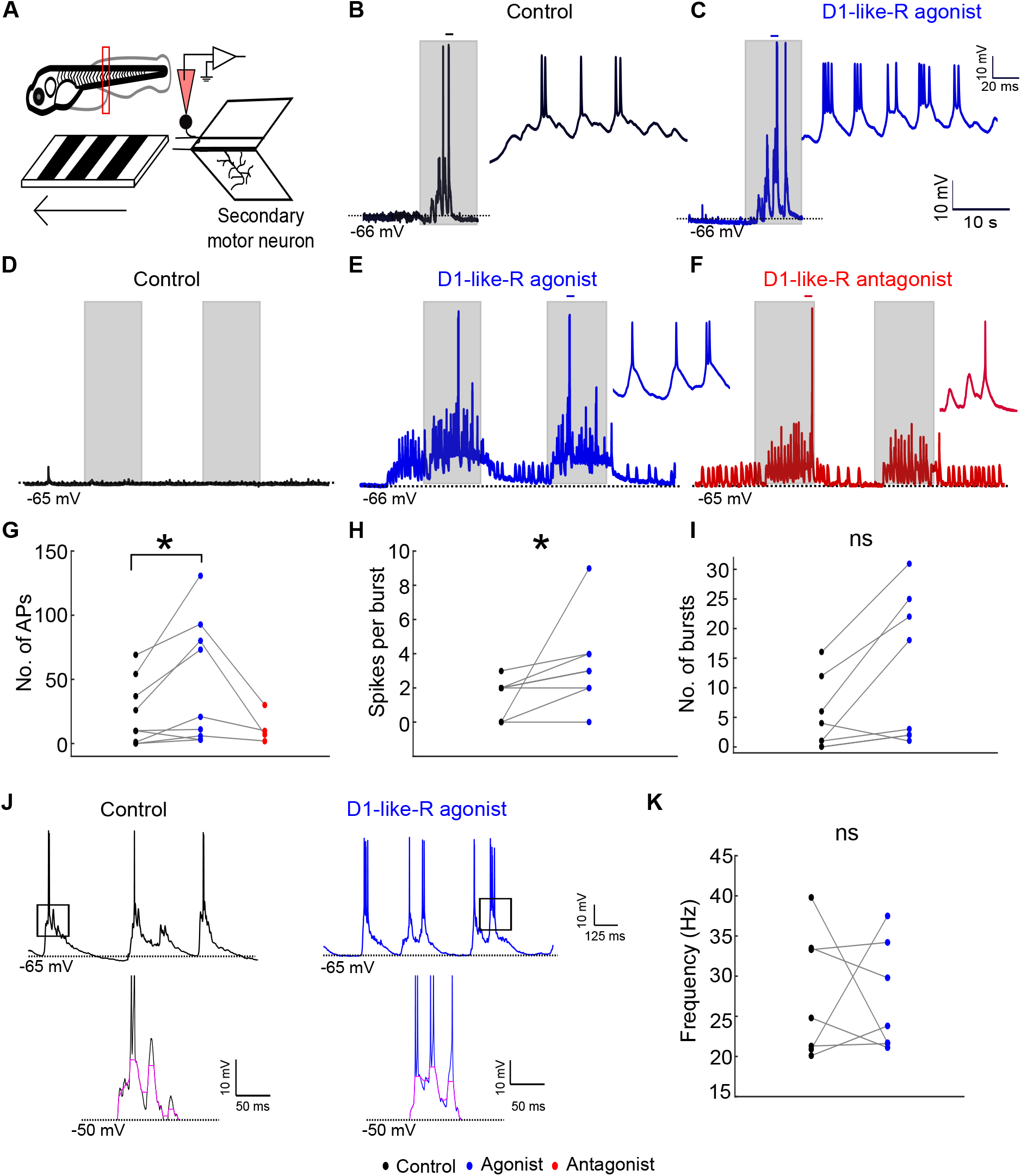
D1-like-R activation enhances drive from secondary motor neurons during OMR. **(A)** Schematic representation of experimental set-up. Whole-cell patch clamp recordings from secondary motor neurons were obtained while presenting 10s of stationary gratings alternating with 10s of forward moving gratings. **(B)** *Left*, recording from a representative secondary motor neuron during a trial. Region under small black line is shown in a zoomed view on the *right*. Shaded areas represent timing of moving gratings presentation. **(C)** *Left*, recording from the same neuron as in **(B)** post application of 20 μM D1-like-R agonist, SKF-38393. Region under the small blue line is shown expanded on the *right***. (D)** Representative recording from a secondary motor neuron that was quiescent during moving gratings presentation before any drug application (black). **(E)** Recording from the same secondary motor neuron as in **(D)** after subsequent application of D1-like-R agonist (blue) and **(F)** D1-like-R antagonist (red). **(G)** Paired plot of number of action potentials fired during moving grating presentation before (black), after application of D1-like-R agonist (blue) and post subsequent application of D1-like-R antagonist (red). **(H)** Paired plot of average number of spikes per burst before (black) and after application of D1-like-R agonist (blue) **(I**) Paired plot of number of bursts during moving gratings presentation before (black) and after (blue) application of D1-like-R agonist, SKF-38393. **(J)** *Top*, representative recordings from a secondary motor neuron during moving grating representation before (black) and after (blue) application of D1-like-R agonist. *Bottom*, zoomed view of region marked with black rectangle showing voltage oscillations (highlighted in pink). **(K)** Average frequency of voltage oscillations before (black) and after (blue) application of D1-like-R agonist. N=9 cells from 9 larvae, Wilcoxon signed-rank test; *p<0.05, ns: not significant. See also Table S1, Figure S1, Figure S2.

Each motor neuron burst has been shown to correspond to motor bursts in fictive swims and is a measure of swim frequency [2, 38]. Further analysis of bursts showed an increase in average number of spikes per burst without a significant increase in the number of bursts post D1-like-R activation (Figure 4H, I). Voltage oscillations underlying these bursts reflect synaptic drive to these motor neurons (Figure 4J; [38]). Average frequency of these oscillations did not differ significantly post application of D1-like-R agonist (Figure 4K). This suggests that upstream rhythm generating circuits are unlikely to be modulated by D1-like-R signaling.

These results indicate that activation of D1-like-R signaling increases swim speed by enhancing drive from secondary motor neurons. This in turn is mediated by recruitment of quiescent neurons and increased firing per swim cycle of already active neurons. These changes point to D1-like-R mediated increase in the excitability of secondary motor neurons, although an increase in synaptic drive during OMR cannot be ruled out.

### D1-like-R activation reduces spike latency and action potential threshold of secondary motor neurons

To test if D1-like-R signaling directly affects the excitability of secondary motor neurons, or if it engages synaptic drive operating during OMR, we asked if secondary motor neurons show changes in firing behavior in response to direct current injections when D1-like R agonist is applied. We analyzed firing responses of secondary motor neurons to a series of depolarizing current steps (25 pA-400pA, 25 pA step size) before and after application of 20 µM D1-like-R agonist (Figure 5A). D1-like-R agonist significantly reduced first spike latency (Figure 5B, C). The decrease in spike latency was observed for all the current steps and was less pronounced when injected currents were high (Figure 5D).

**Figure 5.**
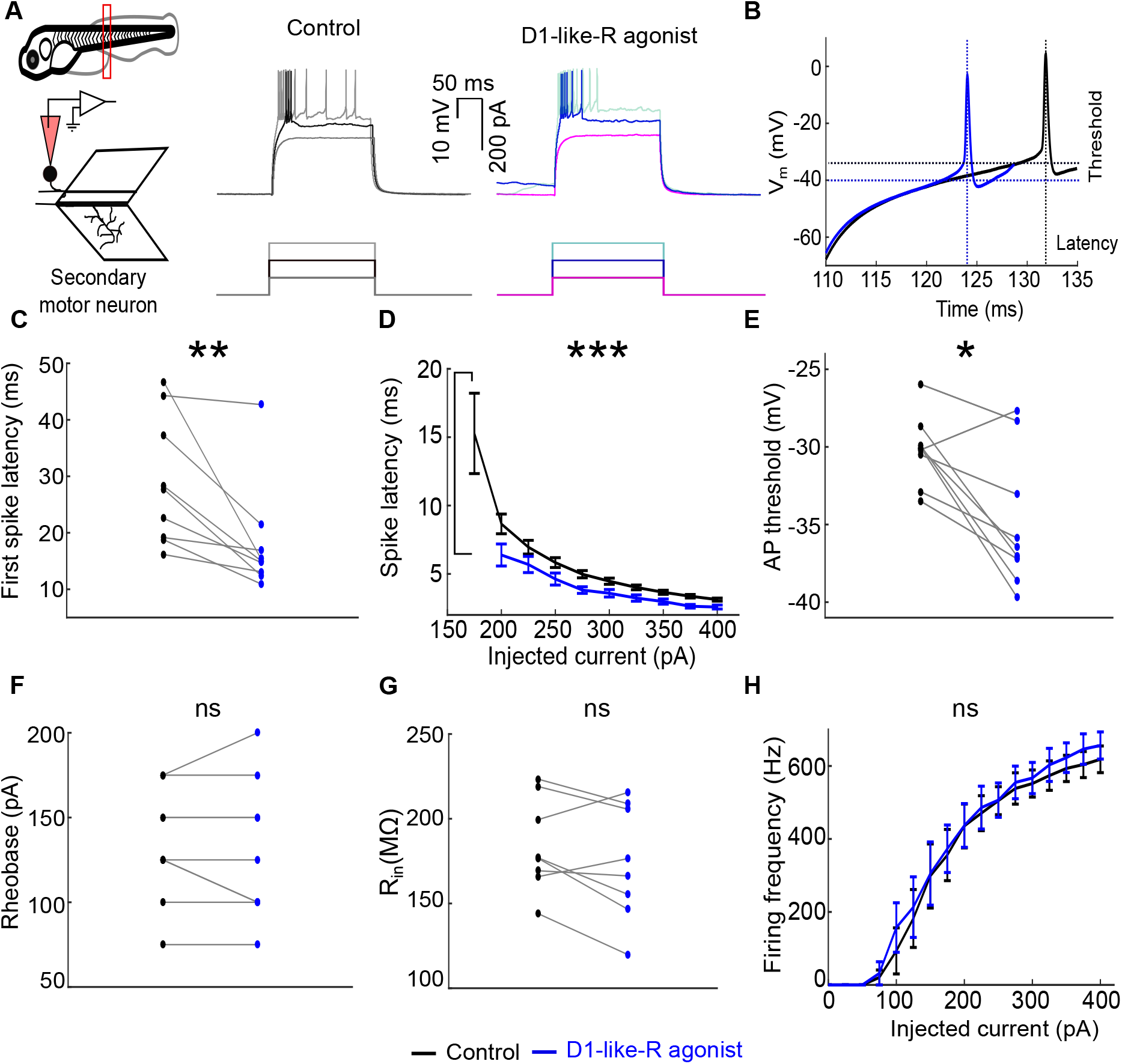
D1-like-R activation reduces spike latency and action potential threshold of secondary motor neurons. **(A)** Schematic of experimental set up. Region marked by the red rectangle is shown below. *Right*, Responses of a secondary motor neuron to a series of depolarizing current steps were compared before (Control) and after application of 20 μM D1-like-R agonist, SKF-38393. **(B)** Representative traces before (black) and after (blue) application of D1-like-R agonist. Vertical dotted lines indicate spike latency and horizontal dotted lines mark threshold for action potential generation before (black) and after (blue) D1-like-R agonist application. **(C)** Summary data for first spike latency. Wilcoxon signed-rank test; **p<0.01 **(D**) Summary data of spike latency for each step of depolarizing current injected. Mixed-Effects model; p<0.0001 **(E)** Summary data for threshold of action potential generation. Wilcoxon signed-rank test; *p<0.05. **(F)** Summary data for rheobase. Paired ttest, p=0.59 **(G)** Summary data for input resistance. Wilcoxon signed-rank test; p= 0.07 **(H)** Instantaneous firing frequency of the first two spikes measured in response to series of current steps before (black) and after (blue) application of D1-like-R agonist. Mixed-Effects model; p= 0.822. ns: not significant. N=9 secondary motor neurons recorded from 9 larvae. Error bars represent SEM. See also Figure S3.

D1-like-R activation also reduced the voltage threshold for the first action potential generated (Figure 5 B, E).

We observed no significant effect of D1-like-R activation on rheobase, input resistance and instantaneous firing rate of first two spikes (Figure 5 F-H). Resting membrane potential, spike afterhyperpolarization (AHP) and spike-width did not change post application of D1-like-R agonist (Figure S3).

These results show that D1-like-R activation increases secondary motoneuronal excitability by reducing spike latency and action potential threshold.

### Quiescent primary motor neurons are recruited during OMR by D1-like-R activation

As primary motor neurons are active only during fast and vigorous swimming [1, 37], we next examined if D1-like-R signaling can modulate primary motor neuron’s excitability as well to bring about faster locomotion. We obtained whole-cell patch clamp recordings from primary motor neurons while presenting drifting gratings from below to evoke OMR (Figure 6A). Primary motor neurons usually showed sub-threshold responses (Figure 6C) or were quiescent (Figure 6F) during visual stimulus presentation. Bath application of 20 µM D1-like-R agonist increased excitability of primary motor neurons. 6/12 primary motor neurons recorded showed recruitment during moving grating presentation post application of D1-like-R agonist (Figure 6D, G). Subsequent application of D1-like-R antagonist on 4 motor neurons reduced the number of action potentials generated in all these cells (Figure 6E, H). Primary motor neurons are further classified as caudal primary (CaP), middle primary (MiP) and rostral primary (RoP) based on the rostrocaudal position of their somata. Of the six primary motor neurons that were recruited, 3 were identified as CaP, 2 as RoP and 1 as MiP. In contrast, number of action potentials spontaneously generated during stationary grating presentation did not differ significantly before and after application of D1-like-R agonist (Figure S2B).

**Figure 6.**
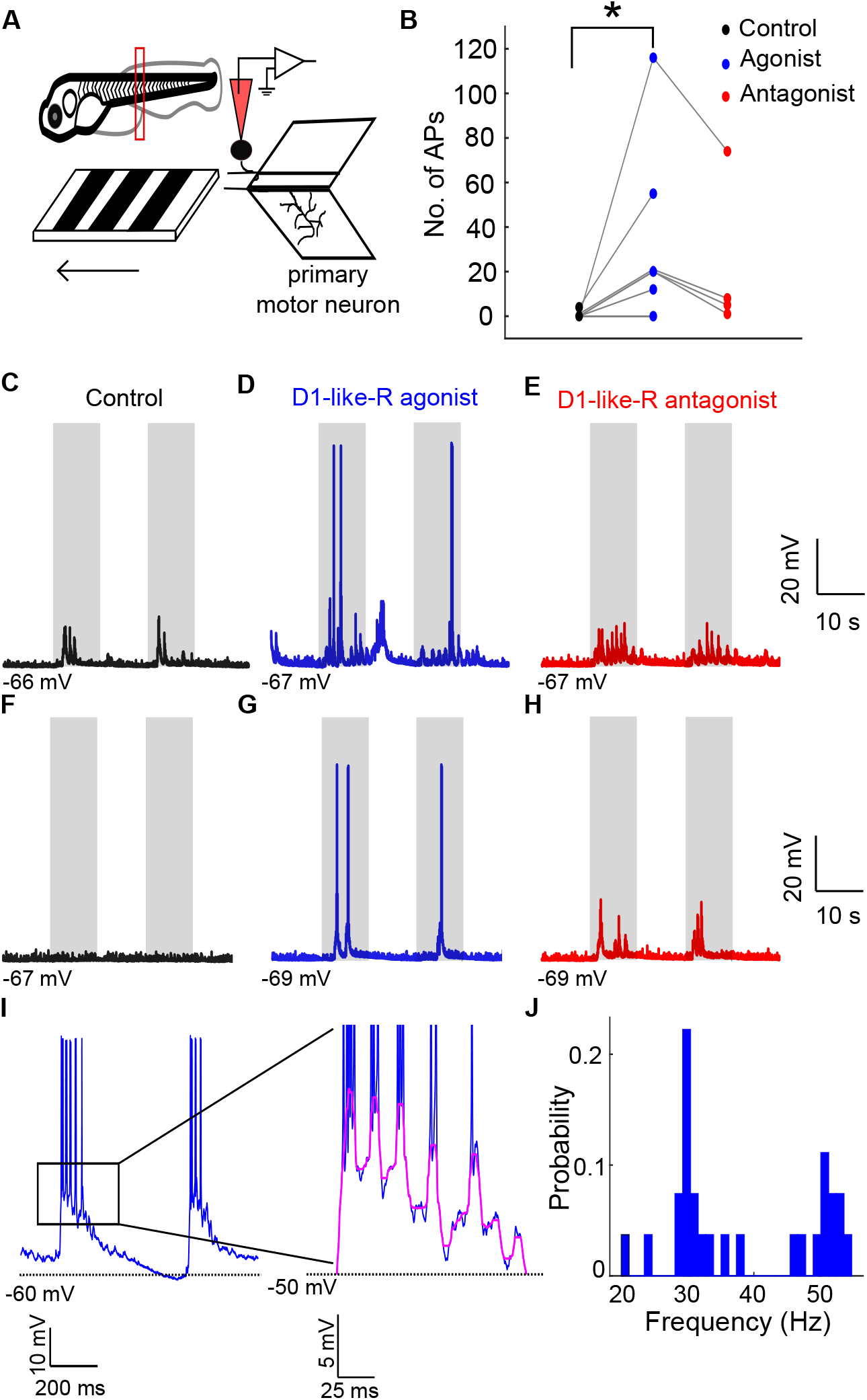
Quiescent primary motor neurons are recruited during OMR by D1-like-R activation. **(A)** Schematic representation of experimental set-up. Whole-cell patch clamp recordings from primary motor neurons were obtained while presenting 10 s of stationary gratings alternating with 10 s of forward moving gratings. Motor neurons were targeted in the region marked by the red rectangle. Zoomed view of the same region is shown on the right. **(B)** Number of action potentials fired by primary motor neurons during forward moving grating presentation before (black), after bath application of 20 μM D1-like-R agonist (blue) followed by bath application of 20 μM D1-like-R antagonist (red) **(C-E)** Representative recordings from a rostral primary motor neuron (RoP) before any drug application (black), after application of D1-like-R agonist (blue) followed by D1-like-R antagonist (red) application. **(F-H)** Recordings from a middle primary motor neuron (MiP) before any drug application (black), after application of D1-like-R agonist (blue) followed by D1-like-R antagonist (red) application. Shaded areas represent timing of moving gratings presentation. **(I)** *Left*, recording from a caudal primary motor neuron (CaP) during forward moving grating presentation post application of D1-like-R agonist. *Right*, zoomed view of the highlighted region showing rhythmic voltage oscillations (marked in pink). **(J)** Distribution of frequency of rhythmic oscillations from all the primary motor neurons that were recruited post application of D1-like-R agonist. N=12 primary motor neurons recorded from 12 larvae, Wilcoxon signed-rank test; *p<0.05. See also Table S1, Figure S1, Figure S2.

Frequency of voltage oscillations underlying bursts (Figure 6I, J) were found to be similar to swimming frequency at which primary motor neuron have been shown to be recruited [1]. These results show that D1-like-R signaling recruits novel neurons from the ‘fast’ motor pool to increase the speed of locomotion during OMR.

### D1-like-R activation increases excitability of primary motor neurons

Next, we investigated the mechanism by which D1-like-R activation recruits primary motor neurons during OMR. Responses to a series of depolarizing current steps were obtained from primary motor neurons. Bath applications of 20 µM D1-like-R agonist were associated with an increase in the number of evoked action potentials (Figure 7A). D1-like-R activation modulated several membrane properties to increase the excitability of primary motor neurons. Spike analysis revealed a decrease in first spike latency (Figure 7B, C). Significant reduction in spike latencies were observed for all current steps injected (Figure 7D).

**Figure 7.**
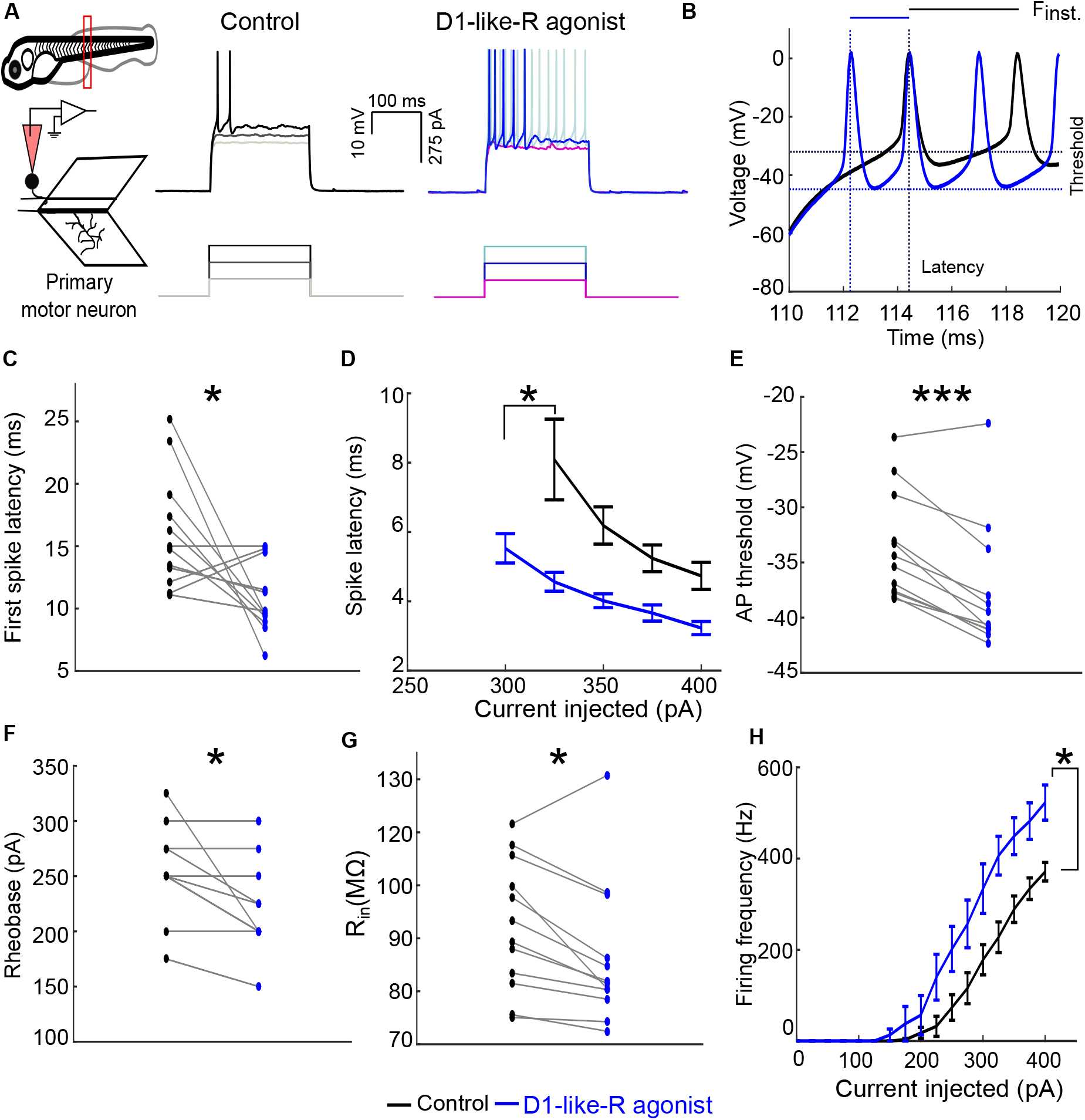
D1-like-R activation increases excitability of primary motor neurons. **(A)** Schematic of experimental set up. Region marked by the red rectangle is shown below. Responses of a representative primary motor neuron to a series of depolarizing current steps were compared before (Control) and after application of 20 μM D1-like-R agonist, SKF-38393. **(B)** Representative traces before (black) and after (blue) application of D1-like-R agonist. Vertical dotted lines indicate spike latency, horizontal dotted lines mark threshold for action potential generation and lines on top indicate first inter-spike interval (F_inst_) before (black) and after (blue) D1-like-R agonist application. **(C)** Summary data for first spike latency. Wilcoxon signed-rank test; *p<0.05 **(D**) Summary data of spike latency for each step of depolarizing current injected. Mixed-Effects model; *p<0.05 **(E)** Summary data for threshold of action potential generation. Wilcoxon signed-rank test; ***p<0.001. **(F)** Summary data for rheobase. Wilcoxon signed-rank test; *p<0.05. **(G)** Summary data for input resistance. Wilcoxon signed-rank test; *p<0.05. **(H)** Instantaneous firing frequency of the first two spikes measured in response to series of current steps before (black) and after (blue) application of D1-like-R agonist. Mixed-Effects model; *p<0.05. N=12 primary motor neurons recorded from 12 larvae. Error bars represent SEM. See also Figure S4.

D1-like-R activation also reduced threshold for action potential generation (Figure 7B, E) and rheobase (Figure 7F). However, a decrease in input resistance was observed post applications of D1-like-R agonist (Figure 7G). Modulation of these membrane properties significantly increased the instantaneous firing rate of first two spikes (Figure 7H), suggesting that with D1-like-R activation, the same synaptic drive to primary motor neurons could still elicit stronger firing. Together, our analysis of secondary and primary motor neuron firing properties in response to grating stimuli and current injections reveal a stimulatory effect of D1-like-R signaling. Activation of D1-like-R increases the number of action potentials fired per burst by decreasing spike latency and spike threshold. The additional spikes fired per burst will act to increase the extent of tail bends, thus increasing the displacement generated in the same amount of time. While not ruling out actions of D1-like-R elsewhere in the motor hierarchy, these effects at the level of motor neurons provide a parsimonious mechanism for the speed regulation observed during OMR.

## Discussion

### Neuromodulatory selection of novel neurons

Zebrafish larvae generate slow swims when presented with slowly drifting gratings. During this behavior they activate secondary motor neurons belonging to the ‘slow’ module, while primary motor neurons belonging to the ‘fast’ module are quiescent. However, when D1-like receptors are activated, primary motor neurons are recruited and secondary motor neurons fire more action potentials, resulting in larger tail bends and therefore greater displacement and faster swims. Our results demonstrate that even under conditions where the sensory input is constant, neuromodulators can remap sensorimotor transformations, such that the behavioral output is altered. In addition, our experiments underline the fact that the pattern of recruitment of neurons during behaviors is heavily dependent on the neuromodulatory context. Recent advances in microscopy and the development of novel neuronal activity sensors have allowed us to map neuronal circuits that are active during a variety of behaviors. Our study serves as a reminder that such maps are a function of the neuromodulatory status of the animal.

### Dopaminergic modulation of motor neuronal excitability

In zebrafish, spinal dopaminergic projections arise solely from the A11 cluster, also known as dopaminergic diencephalospinal neurons (DDNs) [27, 33]. Dopaminergic neuromodulation underlies developmental maturation of swim motor patterns [23] and in particular, the DDNs has been shown to mediate the switch from long to short swim bouts [24]. DDNs are more likely to fire bursts of action potentials during swim bouts and fire tonically during periods of rest [33]. Since burst firing could potentially lead to higher calcium influx into presynaptic terminals, more dopamine is likely to be released by bursting DDNs onto their spinal targets during periods of active locomotion. These findings lend support to our view that endogenous dopamine is capable of altering locomotor speeds *in vivo* in the freely moving animal.

Tyrosine hydroxylase (TH) immunoreactive fibers in the spinal cord make contact with both primary and secondary motor neurons [39] implying direct modulation of motoneuronal excitability by A11 dopaminergic neurons. We show that activation of D1-like receptors enhances the excitability of primary and secondary motor neurons, most likely by acting directly on them. Importantly, we show that both spike latency and threshold are decreased, suggestive of direct dopaminergic modulation of the transient A-type potassium conductance (I_A_). Indeed, dopaminergic reduction of I_A_ leading to a decrease in first spike latency has been reported in invertebrate [40] and vertebrate [41] motor neurons. In addition in primary motor neurons, neuronal gain is increased, suggesting modulatory actions on voltage dependent conductances [42]. These results complement previous studies on mechanistic bases of speed control. The ‘size principle’ states that neurons are sequentially recruited as the speed increases such that the smaller, high input resistance secondary motor neurons get recruited at slow speeds and the larger, low input resistance primary motor neurons are recruited at high speeds. To match this recruitment order, gradations in endogenous rhythmicity have also been observed [38]. In addition, at higher speeds, primary motor neurons preferentially receive enhanced excitation from pre-motor interneurons to enable their recruitment [43]. In contrast, dopaminergic neuromodulation does not seem to tap into any of these previously identified mechanisms of speed regulation. Instead, the neuromodulatory regulation of speed that we identified appears to target intrinsic spike generation mechanisms directly to effect recruitment. Although we show specific modulation of intrinsic properties of motoneurons, modulation of spinal and supra-spinal pre-motor circuits cannot be ruled out.

### Regulation of speed during OMR by spinal interneurons

The recruitment patterns of spinal pre-motor interneurons during OMR has thus far not been documented. Nevertheless, recruitment patterns across a range of locomotor speeds in other contexts has been reported in both larval and adult zebrafish. In adult zebrafish, premotor V2a interneurons connect preferentially to slow, intermediate or fast motor neurons forming distinct speed modules that are sequentially recruited as speed increases [8–10]. However, in larval zebrafish, premotor interneurons active at slow speeds are silenced as swim speed increases, and distinct sets of interneurons are recruited from quiescence at faster speeds [2]. Such a phenomenon has also been reported for speed-related gait changes during limbed locomotion in mice [3–5, 44]. In our studies, the dopaminergic modulation of speed during OMR could be augmented by enhanced suppression of slow-speed interneurons and excitation of fast-speed interneurons. Selective suppression of slow-speed interneurons [45] or activation of fast-speed interneurons [46] are known to play roles in switching swim speeds. Further experiments targeting these interneuronal types should throw light on whether they are targets of dopaminergic neuromodulation.

### Supraspinal circuits governing the OMR behavior

OMR relies on circuits distributed across the nervous system. Retinal input, including responses from direction-selective RGCs, arrives in arborization fields (AF) 5 and 6, which are innervated by the dendrites of pre-tectal neurons. Pretectal neurons show direction-selective tuning, respond to binocular optic flow and send long range projections to the cerebellum and the hindbrain reticular formation [47, 48], in turn leading to the generation of the appropriate swim command in the spinal cord. Several hindbrain neurons are activated when the fish perform OMR [50]. Among these, neurons in the nucleus of the medial longitudinal fasciculus (nMLF) are active during forward OMR and the ventromedial spinal projection neurons (vSPNs) are active during turns [49, 51]. The nMLF makes direct synaptic contacts with primary and secondary motor neurons [52], regulates swim speed during OMR [29] and receives dense innervation from TH-positive fibers [53].

Recently, neurons in the posterior tuberal nucleus (PTN), a sub-group of the A11 dopaminergic cluster, were identified to show large calcium transients in response to forward moving gratings in larval zebrafish [54]. The activation of PTN dopaminergic neurons was delayed by many seconds after optic flow started and lasted for several seconds after it was turned off, suggesting a slow build-up of the modulatory drive. Further, activity in the PTN neurons was suppressed following motor bouts implying a feedback mechanism that limits the dopaminergic modulation. Projections from PTN are restricted to the hindbrain and could potentially be the source of TH-fibers projecting to nMLF. In lamprey, dopamine released from the posterior tuberculum activates reticulospinal cells to increase locomotor output through a D1 receptor-dependent mechanism [55]. Taken together, our results, in the context of the above studies, suggest that during OMR, dopaminergic neurons in the A11 cluster act to increase swim speed via their concomitant modulatory actions at the spinal and supraspinal levels.

## Acknowledgements

The authors would like to thank the following sources of funding support: Wellcome Trust-DBT India Alliance Intermediate and Senior fellowships (VT), Department of Biotechnology (VT), Science and Engineering Research Board, Department of Science and Technology (VT), Department of Atomic Energy (VT), and CSIR-UGC fellowship (UJ). The authors would also like to thank Sriram Narayanan, Pushkar Paranjpe and Dilawar Singh for technical assistance, and T.P. Jagadeesh for maintaining the fish facility.

## Author Contributions

Designed experiments (UJ and VT); Performed experiments (UJ); Analyzed data (UJ and VT); Wrote manuscript (UJ and VT).

## Methods

### EXPERIMENTAL MODEL AND SUBJECT DETAILS

All experiments were approved by the institutional animal ethics committee and the institutional biosafety committee. Experiments were performed on Indian wild type zebrafish (*Danio rerio)* larvae between 6 and 7 days post fertilization (dpf) at room temperature (25-28 °C). Larvae have not undergone sex specification at these stages. Larvae were maintained in 14:10 light-dark cycle at 28 °C in E3 medium (composition in mM: 5 NaCl, 0.17 KCl, 0.33 CaCl_2_, and 0.33 MgSO_4_, pH 7.8).

### METHOD DETAILS

#### Free swimming behavior assay

Zebrafish larvae swam freely in a circular arena 55 mm in diameter. A block of agarose in the centre of the arena restricted swimming in a narrower lane 20 mm in width. Videos were acquired at 120 fps using a CMOS camera (FL3-U3-13-S2M-CS, Point Grey, Richmond, USA) with IR illumination provided from below. A long pass filter (715 nm colored glass filter, Thorlabs, New Jersey, USA) was placed in front of the camera to remove the visual stimulus. Radially spinning square wave gratings were displayed on a screen underneath the arena (Figure 1A). Stimulus presentation was performed using Processing [56]. Acquisition of videos and tracking of larval centroid was performed using Bonsai [57].

Average speed was calculated as total distance travelled in trial duration of 60 s. Bout start and end indices were identified from a plot of instantaneous speed vs time (Figure 1B). Distance travelled within a bout was calculated as bout distance and was divided by duration of respective bout to calculate bout speed. Average of these parameters across all bouts per larva was then calculated to give average bout distance and average bout speed.

#### Head-restrained behavior assay

Larvae were embedded in 1.5% low gelling agarose (Sigma-Aldrich; Missouri, USA). E3 medium was added after the agarose had congealed. Agarose around the tail leaving pectoral fins free was removed for observing tail movements. Optomotor response was evoked by presenting black and white gratings moving with spatial period 10 mm and temporal period 10 mm/s. Visual stimulus was projected on a screen below the larva using a DLP projector (Acer C120; Acer Inc., New Taipei City, Taiwan) after reflection by a cold mirror (FM203; Thorlabs, New Jersey, USA). The arena was IR illuminated from below. Videos were acquired at 500 fps using a high-speed camera (Phantom Miro eX4; Vision research, New Jersey, USA) fitted with a zoom lens (AF Nikkor 60 mm f/2.8D, Nikon, Tokyo, Japan) and infrared pass filter (768 nm, Pixelteq, Florida, USA). Each trial consisted of linearly drifting grating presentation for 12s. Three trials before and four trials after the drug application were performed.

Data were analyzed using custom written scripts in ImageJ, IPython [58] and MATLAB (Mathworks, Natick, USA). Videos were processed in Fiji [59] to obtain larval mid-line. A* search algorithm was used to track tail-tip position using custom written macros in ImageJ. Tail-tip position was used for extracting kinematic parameters. Analysis of kinematic variables was performed offline. Struggles were identified as vigorous body movements with large tail bends and were removed from subsequent analyses. Tracking data was segmented into swim bouts (Figure 2B, left). Amplitude was defined as the maximum (peak-to-trough) tail tip displacement in a bout. Time difference between movement onset and offset of a bout was defined as bout duration (Figure 2B, right). These variables were extracted for individual bouts and average values across all bouts per larva were calculated. Tail beat frequency was defined as the time taken to perform a complete oscillation and was calculated using fast fourier transform. Tail bend angle was calculated using head and tail-tip coordinate. Maximum tail bend angle for each turn was extracted (Figure 3B) and average maximum tail bend angle per larva was calculated.

#### Electrophysiology

Whole-cell patch clamp recordings from motor neurons were obtained while presenting caudal-to-rostral moving gratings on a screen below the stage. Larvae were anesthetized in 0.01% MS-222 (Sigma-Aldrich; Missouri, USA) and transferred to a recording chamber. The larvae were pinned down through notochord using fine tungsten wire (California Fine Wire, CA, USA). The MS-222 was then replaced by external solution (composition in mM: 134 NaCl, 2.9 KCl, 1.2 MgCl_2_, 10 HEPES, 10 glucose, 2.1 CaCl_2_, 0.01 D-tubocurarin; pH 7.8; 290 mOsm) and skin along the tail was carefully removed using forceps (Fine Science Tools, Foster City, USA). Muscles in a hemi-segment (between the 10^th^ and 13^th^ myotomes) were carefully removed to expose the spinal cord. Cells were targeted at 4X magnification with a 60X/1.0 numerical aperture (NA) water-immersion objective (LUMPlanFL) on a compound microscope (BX61WI, Olympus; Tokyo, Japan) with differential interference optics (DIC). Patch pipettes were made using thick walled borosilicate capillaries (1.5 mm OD;0.86 mm ID; Warner Instruments, Hamden, CT, United States) pulled to 1-1.5 μm tip diameter using a Flaming brown P-97 pipette puller (Sutter Instruments, Novato, CA, United States). Pipettes were backfilled with K-gluconate internal solution (composition in mM: 115 K gluconate, 15 KCl, 2 MgCl_2_, 10 HEPES, 10 EGTA, 4 ATP Disodium salt; pH 7.2; 290 mOsm) and typically had resistances between 8 and 10 MΩ. Sulforhodamine was added to the patch internal solution for visualization of cell-morphology and axon tracts.

Motor neurons were targeted blind and classified as primary or secondary post hoc based on their cellular properties (Table S1) and responses to current injections. Primary motor neurons fired tonically whereas the dorsal secondary motor neurons showed chattering response to current injections (Figure S1). Six trials of visual stimulation were presented, with each trial consisting of 10s of stationary gratings alternating with 10s of forward moving gratings. Visual stimuli were displayed on an LCD screen (Waveshare, Hongkong) driven by Raspberry Pi (Model B RASP-PI-3, Raspberry Pi Foundation, Cambridge, UK). Visual stimulus presentation and electrophysiological recordings were triggered simultaneously and acquired using Multiclamp 700B amplifier, Digidata 1440A digitizer and pCLAMP software (Molecular Devices, San Jose, USA). Corrections for bridge balance were applied in current clamp mode. Membrane potentials are reported without correction for liquid junction potential which was measured to be +8 mV for the external and internal solutions mentioned above. The data were low-pass filtered at 2kHz using a Bessel filter and sampled at 20kHz at a gain of 1.

Data were analyzed using Clampfit (Molecular Devices, San Jose, USA) and MATLAB (Mathworks, Natick, USA). Clampfit was used to identify spikes and bursts. To calculate the frequency of oscillations underlying bursts, recordings for trials where forward grating was presented, was first filtered to remove spikes and low frequency periodic depolarization. Frequency with maximum power post fast fourier transform of the filtered recording was used to estimate oscillation frequency for a trial.

To determine intrinsic firing pattern of motor neurons, 200 ms long depolarizing current pulses in the range (25-400 pA, step size 25 pA) were applied in absence of visual stimulus presentation. First spike latency was defined as the time between onset of the current step and the peak of the first spike. Spike threshold was defined for the first spike at rheobase as the membrane potential where the speed of voltage change reaches 5mV/ms. Rheobase was defined as the minimum current required to elicit an action potential. Input resistance was calculated by injecting 200 ms long hyperpolarizing pulses (−100pA to −25 pA).

#### Drugs

Drug concentrations for each experiment was determined in pilot experiments (data not shown). Drugs were dissolved in E3 for behavioral experiments and in external solution for electrophysiological experiments.

### QUANTIFICATION AND STATISTICAL ANALYSIS

Data were tested for normality using one sample Konglomorov-Smirnov test. Paired ttest or Wilcoxon signed rank test were performed for comparisons between the two groups in MATLAB. Mixed effects model using nmle package [60] in R was used to test the effect of drug on latency and firing frequency for different values of current injected. Significance level was defined as 0.05. Details on statistical test, p values and sample size can be found in figure legend.

### KEY RESOURCES TABLE

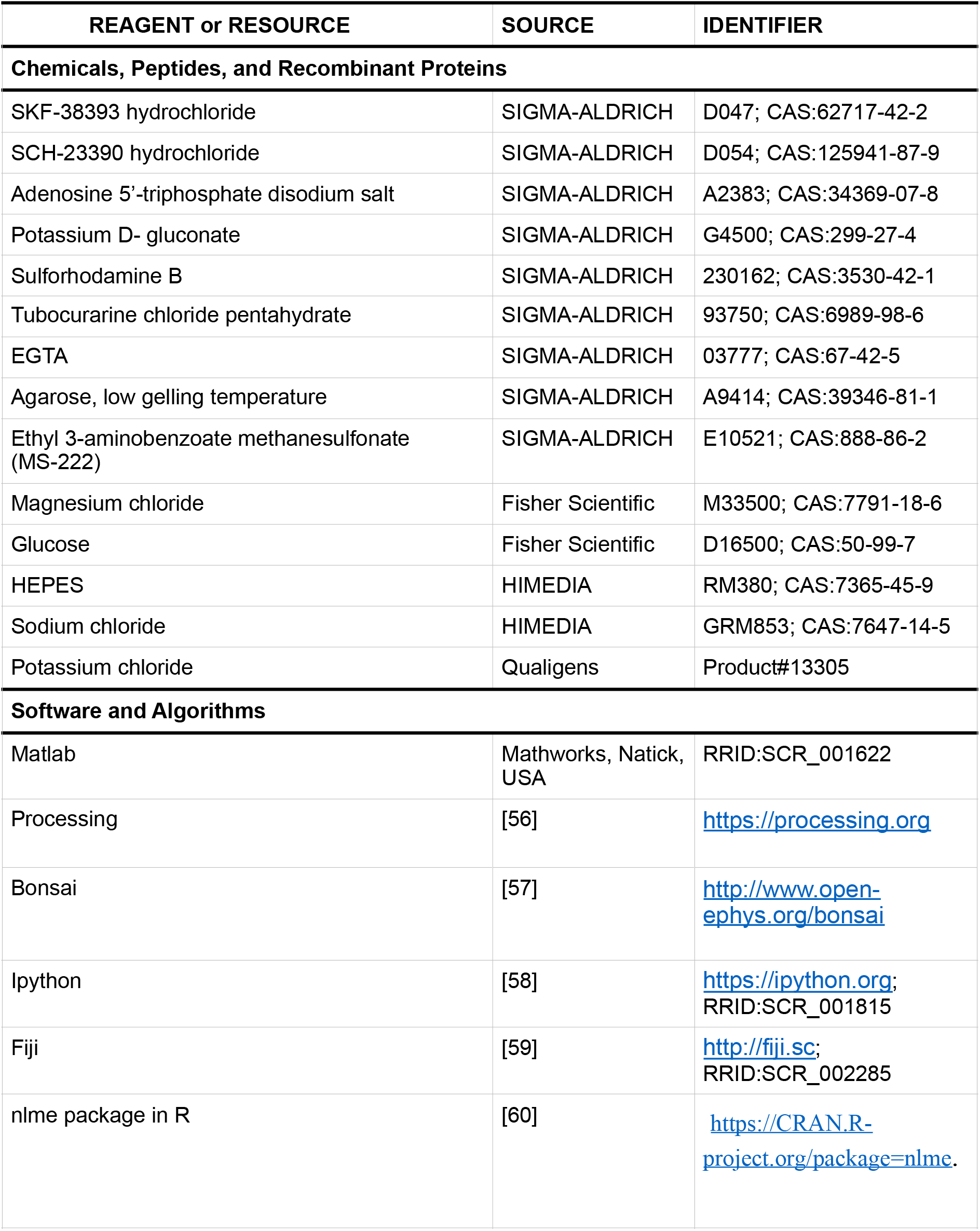

## References

1. McLean, D.L., Fan, J., Higashijima, S., Hale, M.E., and Fetcho, J.R. (2007). A topographic map of recruitment in spinal cord. Nature 446, 71. Available at: https://doi.org/10.1038/nature05588.

2. McLean, D.L., Masino, M.A., Koh, I.Y.Y., Lindquist, W.B., and Fetcho, J.R. (2008). Continuous shifts in the active set of spinal interneurons during changes in locomotor speed. Nat. Neurosci. 11, 1419. Available at: https://doi.org/10.1038/nn.2225.

3. Bellardita, C., and Kiehn, O. (2015). Phenotypic Characterization of Speed-Associated Gait Changes in Mice Reveals Modular Organization of Locomotor Networks. Curr. Biol. 25, 1426–1436. Available at: https://doi.org/10.1016/j.cub.2015.04.005.

4. Gosgnach, S., Lanuza, G.M., Butt, S.J.B., Saueressig, H., Zhang, Y., Velasquez, T., Riethmacher, D., Callaway, E.M., Kiehn, O., and Goulding, M. (2006). V1 spinal neurons regulate the speed of vertebrate locomotor outputs. Nature 440, 215–219. Available at: https://doi.org/10.1038/nature04545.

5. Zhong, G., Sharma, K., and Harris-Warrick, R.M. (2011). Frequency-dependent recruitment of V2a interneurons during fictive locomotion in the mouse spinal cord. Nat. Commun. 2, 274. Available at: https://doi.org/10.1038/ncomms1276.

6. Gabriel, J.P., Ausborn, J., Ampatzis, K., Mahmood, R., Eklöf-Ljunggren, E., and El Manira, A. (2010). Principles governing recruitment of motoneurons during swimming in zebrafish. Nat. Neurosci. 14, 93. Available at: https://doi.org/10.1038/nn.2704.

7. Ampatzis, K., Song, J., Ausborn, J., and El Manira, A. (2013). Pattern of Innervation and Recruitment of Different Classes of Motoneurons in Adult Zebrafish. J. Neurosci. 33, 10875 LP–10886. Available at: http://www.jneurosci.org/content/33/26/10875.abstract.

8. Ausborn, J., Mahmood, R., and El Manira, A. (2012). Decoding the rules of recruitment of excitatory interneurons in the adult zebrafish locomotor network. Proc. Natl. Acad. Sci. 109, E3631 LP–E3639. Available at: http://www.pnas.org/content/109/52/E3631.abstract.

9. Ampatzis, K., Song, J., Ausborn, J., and El Manira, A. (2014). Separate Microcircuit Modules of Distinct V2a Interneurons and Motoneurons Control the Speed of Locomotion. Neuron 83, 934–943. Available at: https://www.sciencedirect.com/science/article/pii/S0896627314006291.

10. Song, J., Dahlberg, E., and El Manira, A. (2018). V2a interneuron diversity tailors spinal circuit organization to control the vigor of locomotor movements. Nat. Commun. 9, 3370. Available at: https://doi.org/10.1038/s41467-018-05827-9.

11. Berg, E.M., Björnfors, E.R., Pallucchi, I., Picton, L.D., and El Manira, A. (2018). Principles Governing Locomotion in Vertebrates: Lessons From Zebrafish. Front. Neural Circuits 12, 73. Available at: https://www.frontiersin.org/article/10.3389/fncir.2018.00073.

12. Kiehn, O. (2016). Decoding the organization of spinal circuits that control locomotion. Nat. Rev. Neurosci. 17, 224–238. Available at: https://www.ncbi.nlm.nih.gov/pubmed/26935168.

13. Arber, S. (2017). Organization and function of neuronal circuits controlling movement. EMBO Mol. Med. 9, 281–284. Available at: https://www.ncbi.nlm.nih.gov/pmc/articles/PMC5331227/.

14. Marder, E., and Thirumalai, V. (2002). Cellular, synaptic and network effects of neuromodulation. Neural Networks 15, 479–493. Available at: http://www.sciencedirect.com/science/article/pii/S0893608002000436.

15. Marder, E. (2012). Neuromodulation of Neuronal Circuits: Back to the Future. Neuron 76, 1–11. Available at: https://www.sciencedirect.com/science/article/pii/S0896627312008173.

16. Harris-Warrick, R.M. (2011). Neuromodulation and flexibility in Central Pattern Generator networks. Curr. Opin. Neurobiol. 21, 685–692. Available at: https://www.ncbi.nlm.nih.gov/pubmed/21646013.

17. Miles, G.B., and Sillar, K.T. (2011). Neuromodulation of Vertebrate Locomotor Control Networks. Physiology 26, 393–411. Available at: https://doi.org/10.1152/physiol.00013.2011.

18. Dickinson, P.S., Mecsas, C., and Marder, E. (1990). Neuropeptide fusion of two motor-pattern generator circuits. Nature 344, 155–158. Available at: https://doi.org/10.1038/344155a0.

19. Lieske, S.P., Thoby-Brisson, M., Telgkamp, P., and Ramirez, J.M. (2000). Reconfiguration of the neural network controlling multiple breathing patterns: eupnea, sighs and gasps. Nat. Neurosci. 3, 600–607. Available at: https://doi.org/10.1038/75776.

20. Sharples, S.A., Koblinger, K., Humphreys, J.M., and Whelan, P.J. (2014). Dopamine: a parallel pathway for the modulation of spinal locomotor networks. Front. Neural Circuits 8, 55. Available at: https://www.ncbi.nlm.nih.gov/pubmed/24982614.

21. Kiehn, O., and Kjaerulff, O. (1996). Spatiotemporal characteristics of 5-HT and dopamine-induced rhythmic hindlimb activity in the in vitro neonatal rat. J. Neurophysiol. 75, 1472–82. Available at: http://www.ncbi.nlm.nih.gov/pubmed/8727391.

22. Barriere, G., Mellen, N., Cazalets, J.-R.J.-R., Barrière, G., Mellen, N., and Cazalets, J.-R.J.-R. (2004). Neuromodulation of the locomotor network by dopamine in the isolated spinal cord of newborn rat. Eur. J. Neurosci. 19, 1325–1335. Available at: https://doi.org/10.1111/j.1460-9568.2004.03210.x.

23. Thirumalai, V., and Cline, H.T. (2008). Endogenous Dopamine Suppresses Initiation of Swimming in Prefeeding Zebrafish Larvae. J. Neurophysiol. 100, 1635–1648. Available at: https://www.ncbi.nlm.nih.gov/pmc/articles/PMC2544474/.

24. Lambert, A.M., Bonkowsky, J.L., and Masino, M.A. (2012). The conserved dopaminergic diencephalospinal tract mediates vertebrate locomotor development in zebrafish larvae. J. Neurosci. 32, 13488–13500. Available at: https://www.ncbi.nlm.nih.gov/pubmed/23015438.

25. Puhl, J.G., and Mesce, K.A. (2008). Dopamine Activates the Motor Pattern for Crawling in the Medicinal Leech. J. Neurosci. 28, 4192 LP–4200. Available at: http://www.ncbi.nlm.nih.gov/pubmed/18417698.

26. Whelan, P., Bonnot, A., and O’Donovan, M.J. (2000). Properties of Rhythmic Activity Generated by the Isolated Spinal Cord of the Neonatal Mouse. J. Neurophysiol. 84, 2821–2833. Available at: https://doi.org/10.1152/jn.2000.84.6.2821.

27. Tay, T.L., Ronneberger, O., Ryu, S., Nitschke, R., and Driever, W. (2011). Comprehensive catecholaminergic projectome analysis reveals single-neuron integration of zebrafish ascending and descending dopaminergic systems. Nat. Commun. 2, 171. Available at: http://dx.doi.org/10.1038/ncomms1171.

28. Portugues, R., and Engert, F. (2009). The neural basis of visual behaviors in the larval zebrafish. Curr. Opin. Neurobiol. 19, 644–647. Available at: https://www.ncbi.nlm.nih.gov/pmc/articles/PMC4524571/.

29. Severi, K.E., Portugues, R., Marques, J.C., O’Malley, D.M., Orger, M.B., and Engert, F. (2014). Neural control and modulation of swimming speed in the larval zebrafish. Neuron 83, 692–707. Available at: https://www.ncbi.nlm.nih.gov/pubmed/25066084.

30. Buss, R.R., and Drapeau, P. (2001). Synaptic Drive to Motoneurons During Fictive Swimming in the Developing Zebrafish. J. Neurophysiol. 86, 197–210. Available at: https://doi.org/10.1152/jn.2001.86.1.197.

31. Irons, T.D., Kelly, P.E., Hunter, D.L., MacPhail, R.C., and Padilla, S. (2013). Acute administration of dopaminergic drugs has differential effects on locomotion in larval zebrafish. Pharmacol. Biochem. Behav. 103, 792–813. Available at: http://www.sciencedirect.com/science/article/pii/S0091305712003346.

32. Souza, B.R., Romano-Silva, M.A., and Tropepe, V. (2011). Dopamine D2 Receptor Activity Modulates Akt Signaling and Alters GABAergic Neuron Development and Motor Behavior in Zebrafish Larvae. J. Neurosci. 31, 5512–5525. Available at: http://www.jneurosci.org/content/31/14/5512.long.

33. Jay, M., De Faveri, F., and McDearmid, J.R. (2015). Firing Dynamics and Modulatory Actions of Supraspinal Dopaminergic Neurons during Zebrafish Locomotor Behavior. Curr. Biol. 25, 435–444. Available at: https://doi.org/10.1016/j.cub.2014.12.033.

34. Green, M.H., Ho, R.K., and Hale, M.E. (2011). Movement and function of the pectoral fins of the larval zebrafish (Danio rerio) during slow swimming. J. Exp. Biol. 214, 3111 LP–3123. Available at: http://jeb.biologists.org/content/214/18/3111.abstract.

35. Bainbridge, R. (1958). The Speed of Swimming of Fish as Related to Size and to the Frequency and Amplitude of the Tail Beat. J. Exp. Biol. 35, 109–133. Available at: http://jeb.biologists.org/content/jexbio/35/1/109.full.pdf.

36. Myers, P.Z. (1985). Spinal motoneurons of the larval zebrafish. J. Comp. Neurol. 236, 555–561. Available at: http://doi.wiley.com/10.1002/cne.902360411.

37. Liu, D.W., and Westerfield, M. (1988). Function of identified motoneurones and co-ordination of primary and secondary motor systems during zebra fish swimming. J. Physiol. 403, 73–89. Available at: https://doi.org/10.1113/jphysiol.1988.sp017239.

38. Menelaou, E., and McLean, D.L. (2012). A Gradient in Endogenous Rhythmicity and Oscillatory Drive Matches Recruitment Order in an Axial Motor Pool. J. Neurosci. 32, 10925 LP–10939. Available at: http://www.jneurosci.org/content/32/32/10925.abstract.

39. McLean, D.L., and Fetcho, J.R. (2004). Relationship of tyrosine hydroxylase and serotonin immunoreactivity to sensorimotor circuitry in larval zebrafish. J. Comp. Neurol. 480, 57–71. Available at: https://doi.org/10.1002/cne.20281.

40. Harris-Warrick, R.M., Coniglio, L.M., Levini, R.M., Gueron, S., and Guckenheimer, J. (1995). Dopamine modulation of two subthreshold currents produces phase shifts in activity of an identified motoneuron. J. Neurophysiol. 74, 1404–1420. Available at: https://doi.org/10.1152/jn.1995.74.4.1404.

41. Han, P., Nakanishi, S.T., Tran, M.A., and Whelan, P.J. (2007). Dopaminergic Modulation of Spinal Neuronal Excitability. J. Neurosci. 27, 13192–13204. Available at: http://www.jneurosci.org/cgi/doi/10.1523/JNEUROSCI.1279-07.2007.

42. Kispersky, T.J., Caplan, J.S., and Marder, E. (2012). Increase in Sodium Conductance Decreases Firing Rate and Gain in Model Neurons. J. Neurosci. 32, 10995 LP–11004. Available at: http://www.jneurosci.org/content/32/32/10995.abstract.

43. Kishore, S., Bagnall, M.W., and McLean, D.L. (2014). Systematic Shifts in the Balance of Excitation and Inhibition Coordinate the Activity of Axial Motor Pools at Different Speeds of Locomotion. J. Neurosci. 34, 14046 LP–14054. Available at: http://www.jneurosci.org/content/34/42/14046.abstract.

44. Talpalar, A.E., Bouvier, J., Borgius, L., Fortin, G., Pierani, A., and Kiehn, O. (2013). Dual-mode operation of neuronal networks involved in left–right alternation. Nature 500, 85–88. Available at: http://www.nature.com/articles/nature12286 [Accessed June 11, 2019].

45. Kimura, Y., and Higashijima, S. (2019). Regulation of locomotor speed and selection of active sets of neurons by V1 neurons. Nat. Commun. 10, 2268. Available at: https://doi.org/10.1038/s41467-019-09871-x.

46. Knafo, S., Fidelin, K., Prendergast, A., Tseng, P.-E.B., Parrin, A., Dickey, C., Böhm, U.L., Figueiredo, S.N., Thouvenin, O., Pascal-Moussellard, H., et al. (2017). Mechanosensory neurons control the timing of spinal microcircuit selection during locomotion. Elife 6, e25260. Available at: https://www.ncbi.nlm.nih.gov/pubmed/28623664.

47. Naumann, E.A., Fitzgerald, J.E., Dunn, T.W., Rihel, J., Sompolinsky, H., and Engert, F. (2016). From Whole-Brain Data to Functional Circuit Models: The Zebrafish Optomotor Response. Cell 167, 947–960.e20. Available at: http://www.sciencedirect.com/science/article/pii/S0092867416314027.

48. Kramer, A., Wu, Y., Baier, H., and Kubo, F. (2019). Neuronal Architecture of a Visual Center that Processes Optic Flow. Neuron. Available at: http://www.sciencedirect.com/science/article/pii/S0896627319303757.

49. Orger, M.B., Kampff, A.R., Severi, K.E., Bollmann, J.H., and Engert, F. (2008). Control of visually guided behavior by distinct populations of spinal projection neurons. Nat. Neurosci. 11, 327. Available at: https://doi.org/10.1038/nn2048.

50. Severi, K.E., Böhm, U.L., and Wyart, C. (2018). Investigation of hindbrain activity during active locomotion reveals inhibitory neurons involved in sensorimotor processing. Sci. Rep. 8, 13615. Available at: https://doi.org/10.1038/s41598-018-31968-4.

51. Huang, K.-H., Ahrens, M.B., Dunn, T.W., and Engert, F. (2013). Spinal Projection Neurons Control Turning Behaviors in Zebrafish. Curr. Biol. 23, 1566–1573. Available at: https://doi.org/10.1016/j.cub.2013.06.044.

52. Wang, W.-C., and McLean, D.L. (2014). Selective responses to tonic descending commands by temporal summation in a spinal motor pool. Neuron 83, 708–721. Available at: https://www.ncbi.nlm.nih.gov/pubmed/25066087.

53. McLean, D.L., and Fetcho, J.R. (2004). Ontogeny and innervation patterns of dopaminergic, noradrenergic, and serotonergic neurons in larval zebrafish. J. Comp. Neurol. 480, 38–56. Available at: https://www.ncbi.nlm.nih.gov/pubmed/15515022.

54. Reinig, S., Driever, W., and Arrenberg, A.B. (2017). The Descending Diencephalic Dopamine System Is Tuned to Sensory Stimuli. Curr. Biol. 27, 318–333. Available at: https://www.sciencedirect.com/science/article/pii/S0960982216314464?via%3Dihub.

55. Ryczko, D., Grätsch, S., Schläger, L., Keuyalian, A., Boukhatem, Z., Garcia, C., Auclair, F., Büschges, A., and Dubuc, R. (2017). Nigral Glutamatergic Neurons Control the Speed of Locomotion. J. Neurosci. 37, 9759 LP–9770. Available at: http://www.ncbi.nlm.nih.gov/pubmed/28924005.

56. Reas, C., and Fry, B. (2006). Processing: programming for the media arts. AI Soc. 20, 526–538. Available at: https://doi.org/10.1007/s00146-006-0050-9.

57. Lopes, G., Bonacchi, N., Frazão, J., Neto, J.P., Atallah, B. V, Soares, S., Moreira, L., Matias, S., Itskov, P.M., Correia, P.A., et al. (2015). Bonsai: an event-based framework for processing and controlling data streams. Front. Neuroinform. 9, 7. Available at: https://www.frontiersin.org/article/10.3389/fninf.2015.00007.

58. Pérez, F., and Granger, B.E. (2007). IPython: A System for Interactive Scientific Computing. Comput. Sci. Eng. 9, 21–29. Available at: https://aip.scitation.org/doi/abs/10.1109/MCSE.2007.53.

59. Schindelin, J., Arganda-Carreras, I., Frise, E., Kaynig, V., Longair, M., Pietzsch, T., Preibisch, S., Rueden, C., Saalfeld, S., Schmid, B., et al. (2012). Fiji: An open-source platform for biological-image analysis. Nat. Methods 9, 676–682. Available at: http://www.nature.com/articles/nmeth.2019.

60. Pinheiro J, Bates D, DebRoy S, Sarkar D, R.C.T. (2019). nlme: Linear and Nonlinear Mixed Effects Models. Available at: https://cran.r-project.org/package=nlme.

