## Supplemental figures for "Neuromodulatory selection of motor neuron recruitment patterns in a visuomotor behavior increases speed"

**Table S1. Cell properties of recorded motor neurons.** Related to Figure 4,5,6,7.

| | Secondary motor neurons | Primary motor neurons |
|---|---|---|
| Input resistance (M Ω) | 197.6±15.8 | 83.8±8.7 |
| Membrane capacitance (pF) | 12.8 ±0.8 | 20.0 ±1.2 |

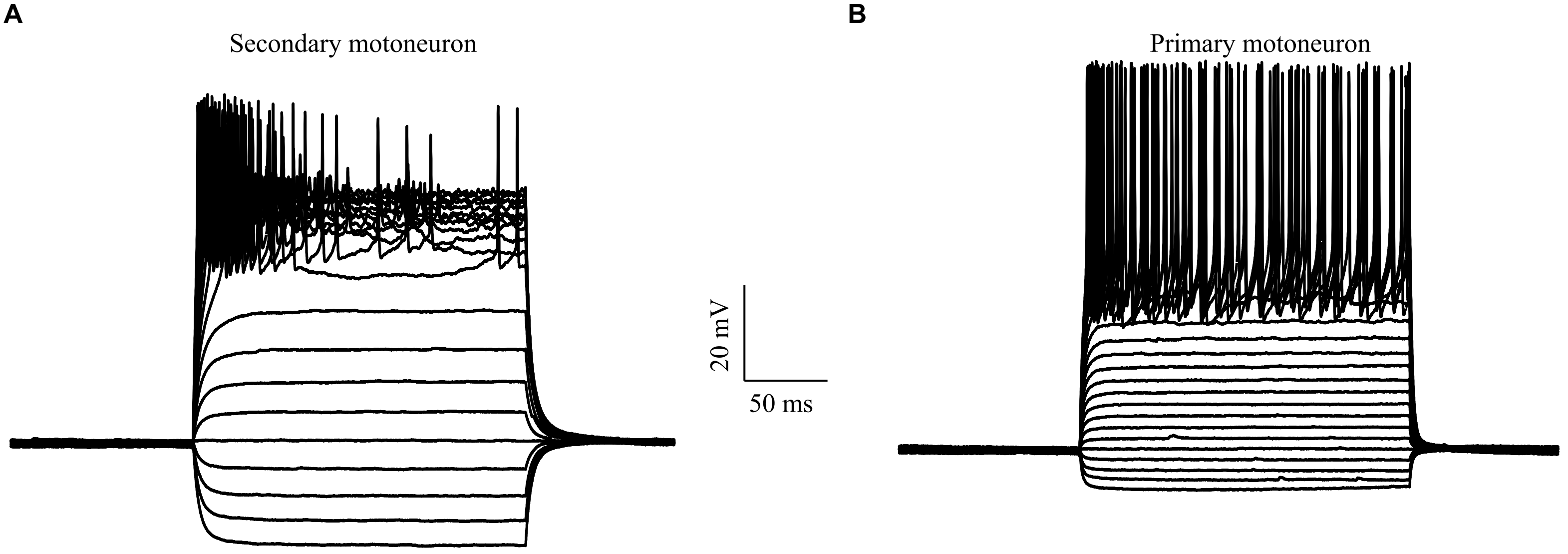


**Figure S1. Characteristic responses of secondary and primary motor neurons to current injections.** Related to Figure 4,5,6,7. **(A)** Chattering reponse of a dorsal secondary motor neuron characterised by intermittent firing. **(B)** Primary motor neurons respond to current steps with stable tonic firing.

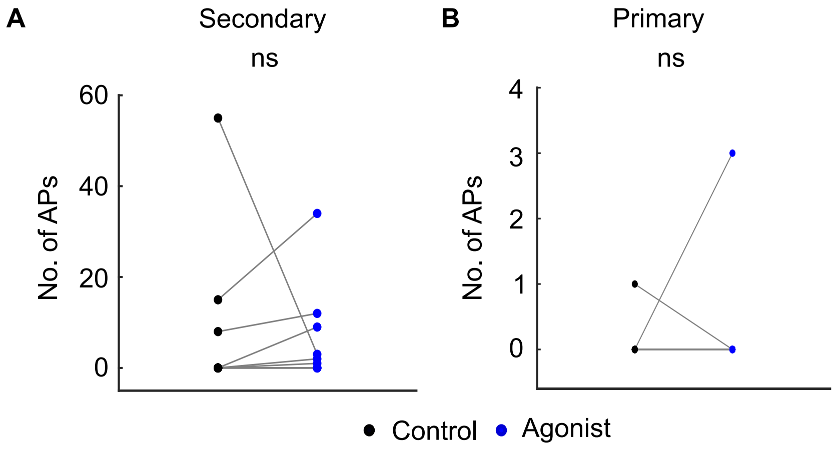


**Figure S2. Modulation of spontaneous action potentials by D1-like-R agonist.** Related to Figure 4,6. Number of action potentials (APs) spontaneously generated during stationary grating presentation before (black) and after (blue) bath application of D1-like-R agonist in **(A)** secondary motor neurons. p=0.4375; Wilcoxon signed-rank test. **(B)** primary motor neurons. p= 1; Wilcoxon signed-rank test. $N_{secondary}$ = 9 cells from 9 larvae, $N_{primary}$= 12 cells from 12 larvae. ns: not significant.

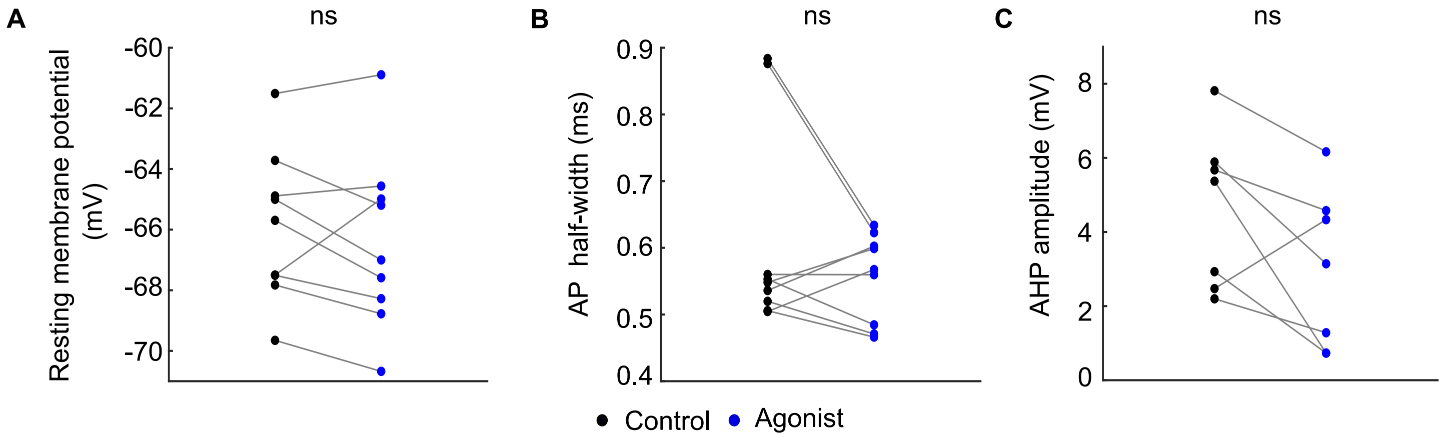


**Figure S3. Modulation of membrane and action potential parameters of secondary motoneurons by D1-like-R agonist.** Related to Figure 5. Paired plot of **(A)** resting membrane potential. p= 0.25; Wilcoxon signed rank test **(B)** action potential half-width. p=0.425; Wilcoxon signed rank test **(C)** afterhyperpolarization amplitude. p=0.1094; Wilcoxon signed rank test. ns: not significant.

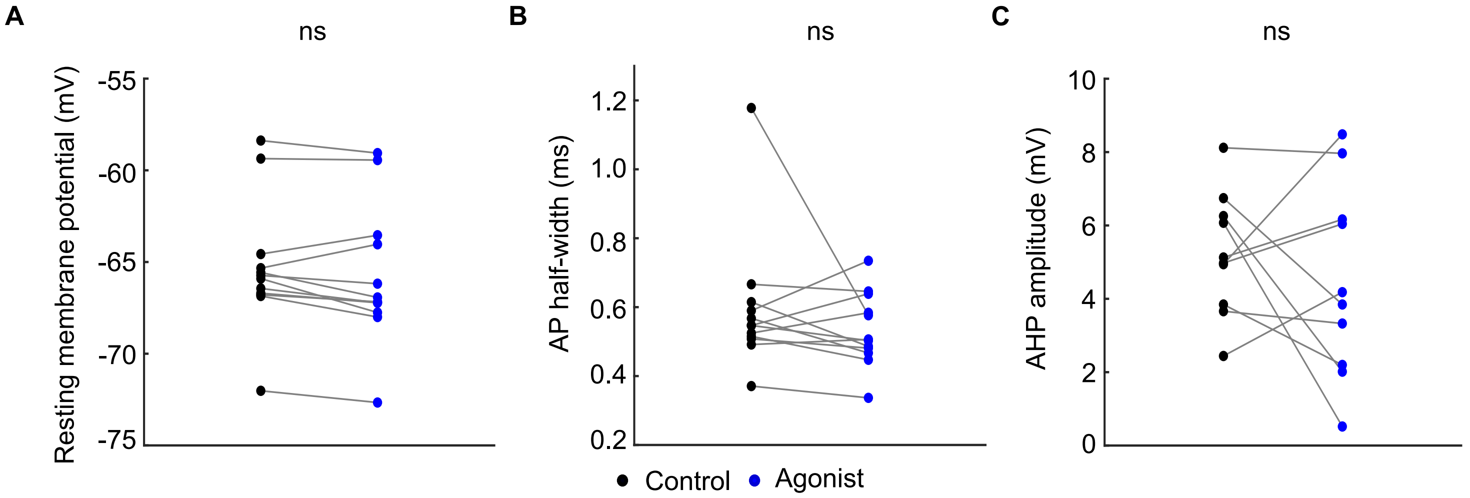


**Figure S4. Modulation of membrane and action potential parameters of primary motoneurons by D1-like-R agonist.** Related to Figure 7. Paired plot of **(A)** resting membrane potential. p=0.1099; Wilcoxon signed rank test **(B)** action potential half-width. p=0.3394 **(C)** afterhyperpolarization amplitude. p=0.4277; ttest . ns: not significant.
